# DN3: An open-source Python library for large-scale raw neurophysiology data assimilation for more flexible and standardized deep learning

**DOI:** 10.1101/2020.12.17.423197

**Authors:** Demetres Kostas, Frank Rudzicz

## Abstract

We propose an open-source Python library, called DN3, designed to accelerate deep learning (DL) analysis with encephalographic data. This library focuses on making experimentation rapid and reproducible and facilitates the integration of both public and private datasets. Furthermore, DN3 is designed in the interest of validating DL processes that include, but are not limited to, classification and regression across many datasets to prove capacity for generalization. We explore the effectiveness of this library by presenting a general scheme for person disambiguation called T-Vectors inspired by speech recognition. These are single vectors created by typically short, though arbitrary in length, electro-encephalographic (EEG) data sequences that uniquely identify users relative to others. T-Vectors were trained by classifying nearly 1000 people using as little as 1 second-long sequences and generalize effectively to users never seen during training. Generalized performance is demonstrated on two commonly used and publicly accessible motor imagery task datasets, which are notorious for intra- and inter-subject signal variability. According to these datasets, subjects can be identified with accuracies as high as 97.7% by simply adopting the label of the nearest neighbouring T-Vectors, with no dependence on task performed and little dependence on recording session, even when sessions are separated by days. Visualization of the T-Vectors from both datasets show no conflation of subjects between datasets, and indicates a T-Vector manifold where subjects cluster well. We first conclude that this is a desirable paradigm shift in EEG-based biometrics and secondly that this manifold deserves further investigation. Our proposed library provides a variety of essential tools that facilitated the development of T-Vectors. The T-vectors codebase serves as a template for future projects using DN3, and we encourage leveraging our provided model for future work.

**Author summary:** We present a new Python library to train deep learning (DL) models with brain data. This library is tailored, but not limited, to developing neural networks for brain-computer-interfaces (BCI) applications. There is abundant interest in leveraging DL in the wider neuroscience community, but we have found current solutions limiting. Furthermore both BCI and DL benefit from benchmarking against multiple datasets and sharing parameters. Our library tries to be accessible to DL novices, yet not limiting to experts, while making experiment configurations more easily shareable and flexible for benchmarking. We demonstrated many of the features of our library by developing a deep neural network capable of disambiguating people from arbitrary lengths of electroencephalography data. We identify a variety of future avenues of study for these representations produced by our network, particularly in biometric applications and addressing the variation in BCI classifier performance. We share our model, library and its associated guides and documentation with the community at large.

## Introduction

Machine learning (ML) has long relied on handfuls of openly accessible datasets, particular to certain sub-fields of study. The MNIST dataset ^1^, for example, has been a common touchstone for image classification and computer vision generally whose use continues (for better or worse) to this day. While a demonstration of state-of-the-art performance on a single dataset, like MNIST, at one time may have been sufficient to warrant the consideration of a new technique, most areas of ML are increasingly comparing performance across *multiple* datasets and tasks. In natural language processing (NLP) for example, the GLUE or SuperGLUE [1] benchmarks express an aggregate score across relevant tasks in the field, varying in size (of dataset) and difficulty. Aspects of the brain-computer interfaces (BCIs) field present a similar but altogether unique variation on this theme. One could loosely divide the work of BCI research (and some of neuroscience more generally) into two streams of inquiry: data *collection* and data *analysis*. Provided perfect – or at least consistent – *collection*, *analysis* techniques are best validated by collecting new data to test hypothetical *analyses*. If *collection* is sufficiently consistent, improvements or subtle changes in *collection* do not strongly impact the interpretation of *analysis* experiments. This is true of other ML fields, but unlike BCI, *collection* in NLP is not a similarly active (or varied) line of research. Instead, quantity of data has taken precedence over quality, or nature [2]. BCI research encourages some specialization in *analysis* or *collection*, with the potential consequence of less-than-ideal *collection* or *analysis* respectively [3]. Likely as a result of both the difficulty in performing *collection* and the need for controlling this confound, thousands of research articles are published per year that solely leverage publicly available BCI datasets [4] which is reminiscent of classical ML. The consequence is that the more general applicability of these results is difficult to judge [5, 6]. The MOABB [4] project is a current effort to establish a wide-ranging benchmark that in no small way addresses this concern, but it remains notably agnostic to *analysis* method, and does not strongly integrate privately collected datasets (see MOABB section).

Intertwined with the desire for stronger generalization claims is the expanded use of DL for BCI classification. While DL is far from established as the preferred *analysis* technique [5], there is undoubtedly keen interest in exploring it [6]. Software solutions for leveraging DL for BCI currently exist (see section on prior work and ecosystem) but, to our knowledge, no publicly accessible DL library is as of yet strongly suited to BCI data, and typically requires significant effort to leverage multiple datasets, public or private. Furthermore, the current ecosystem does not readily support more abstract DL processes, despite that larger *end-to-end* techniques are becoming ever more commonplace in DL [7].

Another concern is that while there are existing *analysis* solutions that employ ML generally, they remain at best simply DL-compatible. This is not necessarily a fault, but it is not uncommon to find DL architectures from the BCI literature that reviews of the literature [5] find poorly motivated or reasoned. It is hard to say precisely why this is, but we speculate that a factor of this phenomenon is that while many BCI practitioners are versed in DL generally, DL remains an enigmatic and fast-moving field, with techniques rapidly falling in and out of fashion. It is here that the question of generalizability returns, in a complementary articulation: *reproducibility*. Often, extreme care is needed to avoid uninformative results due to the notorious and sometimes dramatic dependence on hyperparameters – that often seem innocuous (at first) – to which many DL techniques are prone.

If more generally applicable architectures had a consistent home, and authors wishing to share general models had a semi-standard approach to conform to, a versatile community-driven toolbox of well-motivated techniques could begin to develop and slow the pace of more inexplicable architecture choices. Furthermore, it would allow for stronger benchmarking and provide mechanisms for reproducibility.

Herein we present DN3, the *deep neural networks for neurophysiology* toolbox. This Python library is designed to leverage both public and private data (potentially integrated together) to train deep neural networks in a rapid and reproducible fashion. In particular, we use MNE-Python’s tools for neurophysiological data access [8], storage, and processing, bridged with PyTorch ^2^ – one of the most common and powerful modern deep learning libraries. Knowledge of these underlying libraries is mostly unnecessary, but their lower-level functionalities remains available to DN3. While much of DN3 is tailored to trial-wise BCI classification, it can undoubtedly be used with neurophysiological data more generally. Furthermore, DN3 introduces a unique dataset and experiment configuration tracking module called the `Configuratron`. This module allows datasets to simply reside in the formats they were recorded in, but be automatically prepared for DL processes using short, human-readable descriptions. Then, these descriptions can be easily shared to reproduce work entirely, or simply in design when data cannot be shared. Finally, DN3 implements some existing classifiers and techniques and will remain open source (under a BSD license) to continue to add state-of-the-art techniques to its repertoire, with the goal of remaining convenient for both experts and beginners.

In short, DN3 is best suited to facilitating research at the intersection of deep learning and BCI (and potentially neurophysiological data science more generally, but we focus here on BCI). Experts strongly preferring either of these two fields stand to gain from DN3’s consistency in the abstraction of challenges in the other field. Furthermore, it can dramatically reduce boilerplate code, it introduces general mechanisms for experiment reproducibility, and is fully open-source ^3^.

In section Prior work and ecosystems, we discuss the current Python ecosystem and alternatives to DN3. Next, we provide a structural overview of DN3 and its important modules. Last, we use DN3 to create a cross-paradigm and cross-hardware technique for subject disambiguation (consequently, identification) that we call T-Vectors. We suggest that this technique would have been a long and difficult undertaking but, by making good use of DN3, we have produced a (freely reusable) model with minimal code. T-Vectors are a technique for producing single vectors from variable-length-snippets of EEG data that robustly identify subjects. A neural network was pre-trained to classify over 1000 subjects and subsequently generalized without further training to completely unseen data, despite being recorded in different labs with different hardware. Novel subjects can be identified with well over 90% accuracy using nothing more than the labels of the nearest-neighbouring T-Vectors. We found our representation specifically robust to which task was being performed by the subject and the particular recording session of a sequence of data, yet not confounded by mixing subjects from multiple datasets.

## Prior work and ecosystems

Python has not historically been the standard choice for data analysis in neuroscience. Its large data-science ecosystem and community of open-source, open-access, and community engagement covers a wide array of useful techniques for analysis. MATLAB^4^ instead has been the neuroscience research standard and in fact includes DL tools. However, few if any new DL approaches publish source code with MATLAB, opting instead for typically either Tensorflow or PyTorch (with a community of researchers and hobbyists constantly translating between the two frameworks). Thus, progress towards merging DL with neuroscience is dependent on the MATLAB developers or experts re-implementing entire processes from scratch.

The MNE project, and MNE-Python [8] in particular, is a powerful set of tools for neuroscience data processing, organization, and analysis that further the large ecosystem of Python-based data-science and is a strong alternative to MATLAB. As such, merging MNE-Python with one of these major Python-based DL libraries is a natural solution to studying neuroscience with DL, and is one that has been adopted by prior work in DL with BCI data [9–11]. In the introduction, we discussed a variety of advantages to having a dedicated toolbox for users coming from either of BCI or DL; it is worth highlighting why it is preferable to *not* simply use MNE-Python and a DL library for every DL-neuroscience experiment. MNE-Python’s toolset is very large, and makes few assumptions as to the ultimate application of the data. As such, the efficient development and evaluation of DL processes is nowhere near a *first-class* concern, and can require significant code development for each application, sometimes resulting in code variations just for minor differences in data.

In essence, we observe that Python is the *de facto* choice for DL and that MNE-Python provides many dataset utilities that can facilitate merging DL and neuroscience. However, there is room to add a more application-specific layer on top of MNE-Python to reduce boilerplate code, unify applications, and facilitate novice-level DL.

### The braindecode package

The braindecode Python package is likely the most similar library to DN3 in many respects. Ostensibly, this package provides utilities to train several well known neural network architectures as trial-wise classifiers or regressors of EEG and MEG data. Additionally, it features tools to use a variety of datasets, notably providing a bridge to the datasets featured as part of the MOABB.

The potential advantages braindecode might have over DN3 include the use of the skorch [12] package (itself a layer above PyTorch) rather than PyTorch alone as the DL workhorse. As skorch develops, models prepared for, and any tools added to this (well maintained) library, will likely serve to extend braindecode in a way not true for DN3. That said, we elected to avoid this due to the limitations caused by the fairly reductionist skorch, which enforces training pipelines that might exclude more eccentric approaches that may prove useful in an area that has no apparent standard mechanisms. Consider that the implementation of adversarial architectures in skorch are difficult, and similarly, procedures such as meta-learning, like MAML [13] or REPTILE [14] may not even be possible. Using an adversarial training paradigm has an existing (albeit) small BCI-specific literature [15], while meta-learning is commonly considered for transfer learning problems, itself a keenly sought after methodology for core DL research and BCI [5, 9]. We preferred to err on the side of flexibility in this regard. The braindecode package also provides utilities to enable explainable AI (XAI) techniques (notably from work done by Schirrmeister *et al.* [10]) that can be used as an (albeit rough) attempt at understanding the operations a neural network is performing. While we do not preclude the addition of an XAI module within DN3, we elected to avoid providing any ready-made XAI solutions for the time being, as there seems to be no apparent standards within BCI for this as such, and there is a large risk that if used too liberally, XAI techniques can prove to be very misleading [16].

This being said, DN3 has notable advantages that were (at the time of writing) not available in braindecode, some of which may be difficult to add without significant re-design. The first and perhaps most interesting difference with DN3 is the `Configuratron`, which is a unique addition. Furthermore, the `Dataset` instances that this subsequently constructs has an *application programming interface* (API) that is readily compatible with many other libraries, such as other deep learning libraries like Tensorflow/Keras, and in fact braindecode. Later, we will discuss more detail about this API, but it notably includes a variety of conveniences not similarly provided by braindecode, including methods for accessing data by subject and session, cross validation across subjects, and methods for constructing DL classifiers on the basis of the Dataset instance. Another important point of difference stems from the fact that DL research is dominated by GPUs and other accelerators that run in parallel to CPU operations. It is important to consider how best to leverage idle CPU time. While PyTorch does most of the heavy lifting to facilitate such considerations, DN3 specifies an entire pipeline that modifies data on the CPU while DL proceeds on accelerators. This is in contrast to braindecode, where data transforms are performed before training, and there is a relatively limited data transform/augmentation system.

### MOABB

The mother of all BCI benchmarks (MOABB) project [4] serves as important inspiration for this work. Their stated purpose of consistently validating techniques against public datasets is well aligned with our own. While it would seem clear at the outset that a *benchmark* is very different from a software library, we felt it useful to clearly identify what these are in this case, as the project also includes tools for data analysis and statistical post-analysis. Considering similarities, MOABB is another Python library that also leverages MNE to load, and at various stages represent a BCI (more specifically than DN3) data. Furthermore, they employ a similar set of abstractions around individual sessions, subjects and datasets (each consisting of a number of the former abstractions from left to right). Aside from not being specifically related to DL at all, MOABB makes another fundamentally different (though reasonable for their application) choice to be *strict* on data, but *lenient* on process (e.g. classifier). DN3 relaxes the strictness on data, trying to more effortlessly integrate public and private data, and expand the propagation of relevant information (such as subject id, channel labels, etc.) more extensively through its full pipeline. Ultimately DN3 is not a competitor to MOABB, but is complementary, and shares many of the same goals. DN3 instead focuses on flexibility in leveraged data and specifically facilitates DL-based analysis, whereas MOABB remains agnostic, and less friendly to new datasets (at least in terms of boilerplate code).

## Structural overview

DN3 was organized around two pillars: preparing data and training deep neural network models. DN3 defines an API that connects these pillars through a single recommended point of interface. Figure 1 gives an overview of the major modules of DN3 and which of the two core aspects each interacts with (left or right of the dashed line).

**Fig 1.**
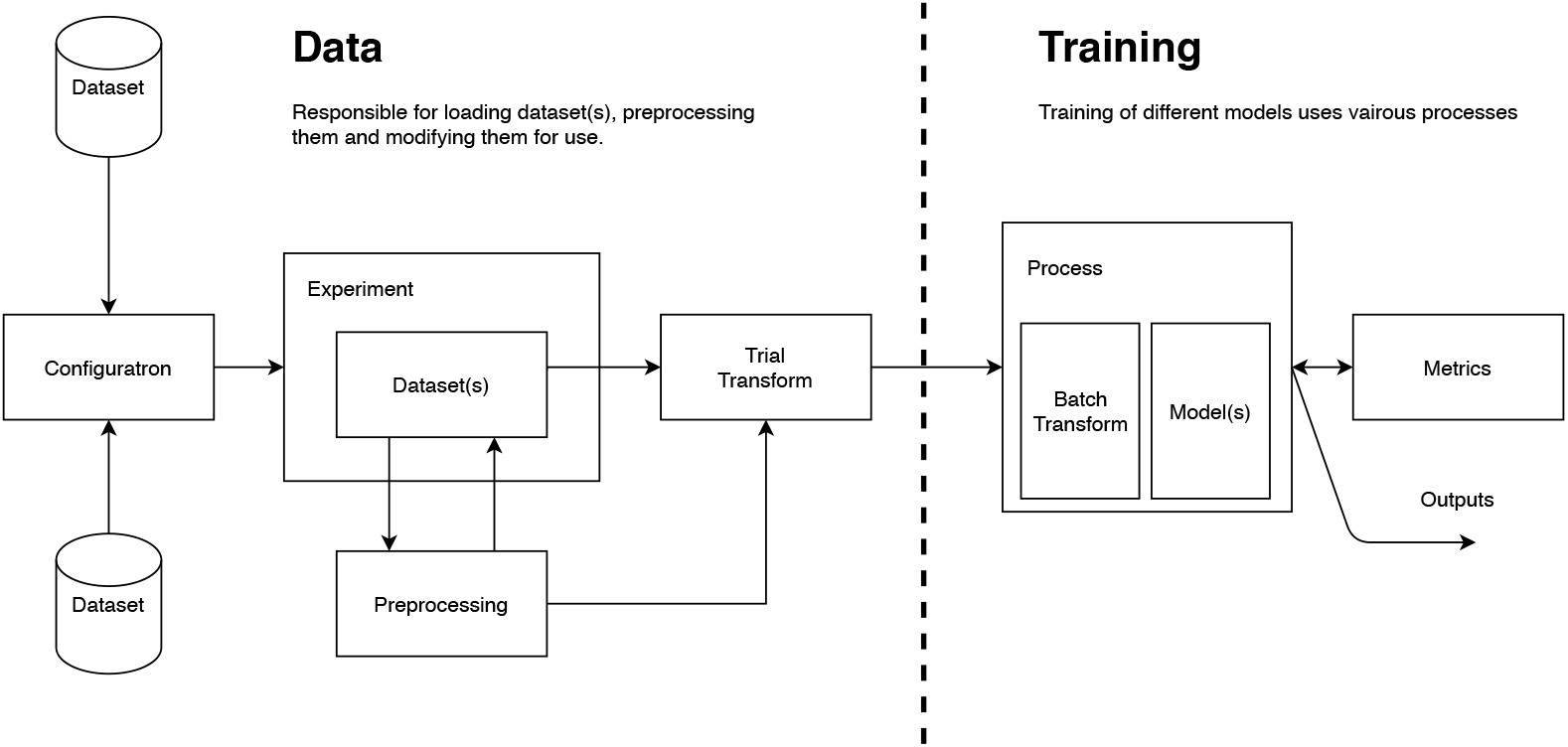
High-level overview of the major aspects of DN3. The library is organized around ease of interface between dataset processing and representation, and the training of deep neural networks with these data.

We elaborate on the motivation of these modules from left to right, focusing on their purposes rather than explaining the particular API which is documented (and kept more up to date than a publication) at https://dn3.readthedocs.io/en/latest/.

### Configuratron

The `Configuratron` is a configuration-file-based dataset organization tool meant to enhance experiment/data preparation consistency and provide significant reductions in boilerplate code. The appropriately formatted configuration files are responsible for listing: the dataset(s) employed in an experiment, where they are stored, whether or what kind of events will be used to crop trials, and more optional pre-processing choices like simple bandpass-filtering or electrode selection. Thus, if a dataset conforms to a consistent directory and naming scheme ^5^ within a common top-level directory, the `Configuratron` easily prepares it for training a DNN simply through a reference to this top-level. From here, splitting the dataset along individual or multiple subjects, and further splitting to the session level is straightforward (see next section).

Some of the more mundane, but highly common, BCI pipeline tasks, such as: renaming or remaping channels, excluding inconsistent or bad subjects and sessions (which in a small capacity is also automatically discovered while loading datasets), and adding some basic transforms such as normalization, are all specified using these configuration files. The majority of these options are meant to provide a much more efficient way of performing the myriad housekeeping (not really even preprocessing) steps that are, for the most part, handled by MNE-Python, but must be done consistently for every subject and session (and dataset if experimenting across datasets). Sharing these configuration files, allows for data to be loaded in, again, a *consistent* fashion between different researchers, encouraging reproducibility but also flexibility as to how the analysis aspect will be performed, without ever exchanging raw/processed data or a complete codebase, which itself may be impossible to share, e.g. due to privacy concerns.

While leveraging standard directory structures and file-types makes for minimal coding, this structure can also be adapted to more custom-solutions. Data can be forcibly injected at the session (see figure 2), person and dataset level, while still leveraging the remaining configuration options when appropriate. Furthermore, the configuration file behaves as a more general place for logging hyperparameters, by allowing arbitrary configuration elements. These can include preprocessing and transform details, or DL hyperparameters. Finally, a simple import system (accomplished through a YAML directive) allows for importing other configurations to the experiment. This allows selecting to import different hyperparameter sets, or pulling hyperparameters from web-enabled hyperparameter search tools.

**Fig 2.**
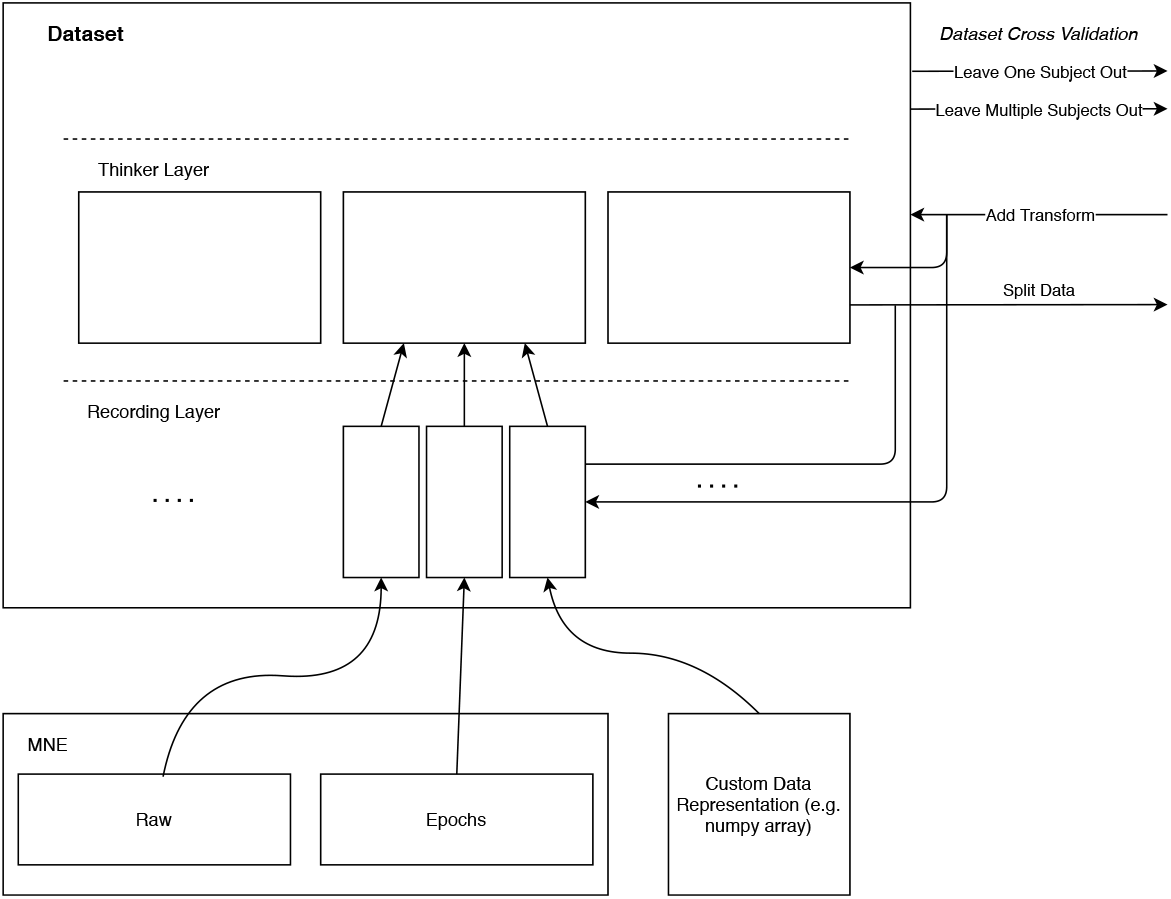
Here the hierarchical structure of a `Dataset` is outlined, with individual sessions made up of the MNE-Python `Raw` and `Epoch` instances, or custom representations for more unique problems. The arrows to the right indicate some common API endpoints for developing training, validation and test sets from the `Dataset`.

### Data

This module implements the higher-level containers that represent a dataset and its constituent parts. `Datasets` are specifically comprised of a set of `Thinkers` that represent each respective subject, and those are each comprised of a number of recordings. Figure 2 further illustrates this hierarchy and shows how MNE `Raw` and `Epochs` objects ^6^ underpin the data layer below these. Thus, while MNE compatible data is naturally integrated into this scheme, `Recording-` and `Thinker`-level APIs allow for more customized data integration into the `Dataset` format.

Once data is represented as a `Dataset`, single data instances are fetched from disk or system memory depending on configuration (important for more large-scale datasets that are beginning to develop [17]). This is combined with *experiment*-focused conveniences like leave one (or multiple) subject(s) out cross-validation, or randomized splits, and also the collection of dataset level data distributions and statistics. Furthermore, DN3 includes a suite of utilities that help develop automated rejection of trials, or other steps to filter for the most pertinent sections of data.

### Preprocessing & Transforms

Commmon to all BCI analysis is some preprocessing step(s), or adjustment of the raw data. For example simply excluding non-neural recording channels, leveraging those channels to strip away artifacts, or spatial filtering/channel re-weighting. DN3 tries to minimize the time spent doing steps like these *before* each experiment by creating a pipeline to transform data while it is loaded. This could be as simple as stripping electro-oculogram (EOG) channels while trials are fetched from disk, though this alone does not really require a pipeline as such. If alternatively, independent component analysis (ICA) signals are to be removed based on correlation to EOG signals, a preprocessing stage would calculate the forward and backward transformation (optionally saved offline), and then the rejection of these components can happen as a batch is collected for DL. These are relatively standard methods, but we created a notably flexible pipeline to avoid precluding things like mixing of transformed and un-transformed trials, mixing different transformations, or doing something more exceptional like converting trials to their T-Vector representations.

### Processes & Trainables

Rather than limiting analysis to classification or regression using neural network models (or set of modules that constitute them), DN3 employs a more abstract training `Processes` API to train… `Trainables`. The distinction may not be clear from the outset Undoubtedly, the `StandardClassification` process is likely to be used the majority of the time, but DL as simply a process of classification or regression with a single loss function is perhaps at the time of writing, more than ever, not sufficient. There are now wide variety of end-to-end systems being developed in DL [14, 18–20] that can not be fully characterized within such a frame. A community driven solution like DN3 could help make more of these available for wider use. Thus, a `Process` in DN3, is a more general formulation than simply classification or regression, and is designed to support diverse uses of backpropagation with respect to a (or even multiple) loss function(s). There are some added assumptions that incoming data is, in a sense, neurophysiological data (i.e. made up of sequence of sampled channels). While conveniences like loading batched (and batch-wise transformed) data to the appropriate accelerator, calculating training and validation metrics and under/over sampling are all readily available through a change in arguments during `Process` creation. In other words, a `Process` encapsulates the procedure by which data is fed to a model and the backpropagation of losses through it. Those that are unconcerned with exactly how particular weights get updated can safely ignore this latter detail, while those that are concerned with questions of why, how, when and if certain weights are changed, can create their own processes.

`Trainables` are the neural network modules, or more generally, modules that integrate into the forward and backward passes in DL. These can simply be PyTorch `Modules` and `Functions`, or the extended wrapper provided by DN3: `DN3BaseModel`(s). The advantage of using the DN3 API, is that models can be generated based on `Dataset` instances, meaning according to the incoming channel set, sequence length, sampling frequency and classification targets. This has been, in our experience, a way to alleviate a major source of tiresome code, particularly in projects that leveraged several datasets (like the one we present below).

Processes would not be complete without the addition of `Metrics` to evaluate the suitability of `Trainables`. To this end we have taken advantage of the existing Python data science ecosystem, and provide utilities to leverage the many metrics provided by standard choices like `sklearn` [21] through wrapper functions and decorators.

## T-Vectors

In this section, we present original work that showcases some of the unique advantages of DN3. We showcase integrating multiple datasets, BCI-dataset-specific utilities, and the capitalization of the wider Python ecosystem.

### Motivation

The recorded features of most subjects performing BCI tasks can vary dramatically both within and between subjects, making universally applicable classifiers difficult to develop [9, 22–24]. The sensory motor rhythm (SMR) paradigm in particular exhibits a form of subject-dependent variability that manifests in some 20-30% of people seemingly incapable of even eliciting expected features for classification in the first place [22, 23]. The nature of this dramatic variability is a topic of continual research but not nearly fully characterized. Prior work groups this variability into psychological, physiological, anatomical, and demographic differences, though integrated studies of these factors are notably lacking [23]. These are then perhaps overly inclusive categories, capturing all known correlates to performance. While these factors potentially inform the cause of the signal variations, the goal nonetheless remains that new users (or new sessions of existing users) can quickly use BCI systems [5, 25, 26]. To this end, previous work has found that identifying *similar* users implies what configuration of features and/or classifiers may be effective for a new context [26]. In other words, transfer learning (TL) can be enhanced by identifying similar enough users.

More than just identifying similar users, Riemannian geometry-based covariance matrix classification is a large step towards better TL in BCI [5, 27]. This alternative represents trials using the covariance matrix of the channels of each trial, resulting in a symmetric positive definite matrix (SPD) confined to its associated differentiable manifold *M* (i.e., the manifold of the set of SPD matrices). The advantage of this approach is the robustness of a trial’s representation on *M* under a variety of conditions, notably under a variety of BCI preprocessing and recording techniques [5, 27]. Specifically, this means that these robust representations allow for better subject-to-subject transfer and stability across different data collection efforts [5]. Prior work has shown that trials of particular users tend to cluster along *M*, and transfer learning can be further improved if these representations are shifted along *M* to a common centralized location [28]; however, this requires the use of Riemannian geometry to establish a central location along *M*, which is cumbersome. Thus the stability afforded by confining trials to the invariant space *M* is traded-off against the limitation in features asserted by the SPD representation and the inconvenience of the geometry, where only some classification algorithms naturally extend.

In the work we present below, we have first and foremost asked whether a deep neural network can be used to create a robust (invariant) descriptive vector of any EEG user. Ideally, this vector could identify a particular user juxtaposed to others, (in the best case) irrespective of hardware used or activity performed. This should consequently then be a model of latent subject-wise variation, where features or directions in this space may be descriptive of subject-wise differences. This is an *indirect*, and data-driven modelling of latent differences, rather than an explicit account of these differences. Towards application, this representation could be immediately leveraged to identify similar subjects and as noted above may be informative of how best to classify their features. However, it presents a variety of additional opportunities. As a consequence, the range of these representations creates a manifold, somewhat like *M* above, but without the constraints of solely covariance-based features and the requirement to use Riemannian geometry (the more standard Euclidean sense should suffice). Evaluating the result of all of these motivations is beyond the scope of this work alone, and here we have focused on the development of a robust learned vector first.

To develop our vectors, we took inspiration from a similar problem that arises in automatic speech recognition (ASR). Unlike BCIs, variability in speech is intuitively apparent – different individuals have high or low-pitched voices, an individual may be yelling, whispering, or attenuating their speech production, or may have a regional accent. This latter case can loosely serve as an analogy for part of the BCI-specific variance problem, if we assumed that non-native speakers leverage co-articulation patterns or colloquialisms from their native tongue when speaking a non-native language, it results in (what we commonly assume to be) the same semantics with a different set of (acoustic) manifestations. This *same semantics, different manifestations* framing is a loose lens that we extend to BCI subject variation (although what a non-native, versus native *speaker* means for BCI strains the analogy). Therefore, this lens was useful insofar as it suggested a possibly effective method to deal with the known phenomenon of person-wise variation, one that was readily transferable to BCI data: summarize utterances into a fixed-length representation that is able to classify a large set of users. I-Vectors and their variants [29] are historically effective, but with more speech recognition being performed by DNNs, the similar X-Vector approach is common and seemingly more accurate [29].

Thus, herein we investigate whether DNN-derived X-Vectors, used mostly as they were presented in the ASR literature, are capable of modelling *thinker* rather than *speaker* instance differences. We call these vectors trained to identify *thinkers*:

### T-Vectors

We would be remiss to not also mention that, while our own motivation comes from a desire to model the latent space for inter-subject variance, our proposed solution (if effective) is also very relevant to a body of research in EEG-based biometrics [30, 31]. Here, EEG signals are leveraged to uniquely identify users, typically in the interest of security applications. Most commonly, this is a problem framed in terms of *authentication* or *identification* [30]. Authentication strives to identify a single person from an open set (and is ostensibly more desirable [30]), while identification considers options in a closed set of candidates. We suggest that T-Vectors represent a *disambiguation* paradigm instead. T-Vectors should disambiguate *any* subject (whether seen during training or otherwise) by (Euclidean) proximity to other vectors. It could thus be adapted to either authentication or identification, but is not immediately one or the other.

### Datasets

An ideal *training* dataset here is one that is as general and representative as possible (while remaining practically tractable) so that it transfers well to many applications. In this case then, it specifically should represent a multitude of different people, performing various tasks, across many demographics and recording contexts, preferably separated in time. To our knowledge, the closest-fitting publicly-available project was the Temple University Hospital EEG Corpus (TUEG) which advertises data from individuals as young as less than a year old, to over 90, a split of 51% female, and sessions that were separated up to eight months apart [17]. It consists of clinical EEG recordings from Temple University Hospital using a limited variety of monopolar EEG systems, all with a roughly 10/20 electrode configuration, in addition to a variety of auxiliary channels including eletro-ocular (EOG), cardio and myographic recordings. The majority of the recordings were made with sampling frequencies of 250 H*z* or 256 H*z*, though some were higher (typically multiples of this, such as 512 H*z* or 1024 H*z*) [17]. We limited our analysis to the version 1.2.0 subset of TUEG, which after rejecting outlying data (see Preprocessing), featured 1364 people ^7^ roughly an order of magnitude larger than any prior work in EEG user identification that we were aware of [30, 31].

For downstream subject disambiguation (datasets considered once the T-Vector model was determined), we focused on two SMR datasets, due to the task’s known intra- and inter-subject variability. First, the well known BCI competition IV, dataset 2a (BCIC) [32], which we selected for its clear isolation of two distinct recording sessions separated by days, and its limited subject set. The limited subject set was particularly valuable for visual examination. It similarly featured a largely 10/20 monopolar recording setup of 22 EEG channels, 3 EOG channels and a single event trigger channel (the EOG and of course trigger channels were simply discarded in this work). Secondly, we used the easily accessible movement and motor imagery database (MMI) [33, 34]. This featured 109 subjects using 64 channels in a 10/10 channel configuration, sampled ostensibly at 160 Hz. Four of these subjects were inconsistently sampled and excluded from training ^8^ leaving 105 possible subjects. Note that these datasets also featured (particularly MMI) in prior work in subject identification for biometric applications [30, 31].

All three datasets were simply downloaded from publicly available locations: TUEG^9^, MMI ^10^ and BCIC ^11^. The `Configuratron` automatically prepared each of the datasets using the configuration files available with the project. This included renaming of a variety of channels, notably from TUEG, excluding a number of uninformative sections of data and more. The most consequential of these is presented in the following section, and can otherwise be found in the published source code.

### Preprocessing

The TUEG dataset contains recordings at various sampling frequencies, all with a low-pass filter that did not violate the Nyquist criterion for a 256 H*z* sampling frequency (i.e., low-pass filter ≤ 128 H*z*) were kept and resampled to 256 H*z*. This resampling was kept as simple as possible: in most cases, a decimating procedure (taking every *n* samples) whenever the sampling frequency was whole multiple of 256 H*z*. Further, any data sampled at 250 H*z* was resampled to 256 H*z* using the nearest sample in time to the original sequence. Note that both the decimation (when sampling frequencies are whole multiples of a standard sampling frequency) and the nearest sample interpolation are done automatically by DN3 when a global sampling frequency is specified for the experiment (a flag set simply in a configuration file). A consistent ordering for the channels was then determined, so that the same EEG channel was consistently found at the same tensor index, e.g. the FP1 channel was always found at index 1. This was necessary to allow the use of data across different recording hardware, and has the benefit of allowing previously trained models to be transferable to other applications. DN3 has a system for doing this automatically called the `Deep1010` mapping, which consistently maps 77 EEG channels, 2 earlobe reference electrodes (this covers the very common 10/20 channel scheme and additionally adds the mid-way points to mostly cover the 10/10 extension [35]) and 11 auxiliary channels that includes 4 electro-oculogram channels (horizontal and vertical for left and right eyes) and 7 miscellaneous channels that allow for the integration of other sources for potential artifacts such as electro-cardio and myograms, additional reference channels or other more application-specific channels. Ultimately, when training and evaluating T-Vectors, all but the EEG channels were removed. We explain the `Deep1010` as it is a unique feature of DN3.

We cropped non-overlapping and reasonably dissimilar sequences by taking 1280 samples (at a sampling frequency of 256 H*z* corresponding to 5 seconds) every 7680 samples (30 seconds). In other words, the first 5 seconds of every 30 from each recording were considered for training. Herein these cropped sequences are referred to simply as *points*. Due to the extreme size of the TUEG dataset, it was difficult to triage the entire dataset, but a cursory look found notable artifacts and many cases of channels with absent (zero, or minimally varying) data. Scaling each point to lie between −1 and 1 ^12^, we considered the overall distribution of these points and determined upper and lower bounds on standard deviation for viable training points. This is described in more detail in appendix A, but it suffices to say that a very significant proportion of data had curiously low or high variation as compared to more controlled/smaller scale datasets, and these extremes were removed. We employed a variety of readily available tools from the DN3 library to accomplish this.

Subsequent to these initial preprocessing steps, each person had, on average, 234 viable points for training, with an overall minimum of 1 and a maximum of 2621, totalling just under 320 000 points.

The BCIC dataset was prepared as if for classification of the associated 4-way SMR task (imagined left and right hand movement, foot movement and tongue movement) [32]. Points were determined by taking 4.5 second crops from 0.5 seconds before the event marker until 4 seconds after, this time period has been shown in prior work to maximize the event-related signal for previous neural network classifiers [9, 10]. Similar to some of the TUEG data, the sampling frequency of 250 H*z* was upsampled using the nearest point in time to the common 256 H*z*

The MMI dataset was prepared differently, focusing instead on leveraging the totality of data available for all 105 subjects. This meant including all 14 sessions performed by each person, and using all of the recorded data. To do this, the sampling frequency of 160 H*z* was adjusted to the 256 H*z* mark again by simply using nearest neighbours upsampling. We note here that while this is nowhere near an ideal choice, we focused on making simple choices to determine how tolerant the T-Vector extraction was. With a unified sampling frequency, each point was made from each non-overlapping 4 second crop of data. The one exception to this was that when investigating the capacity of T-Vectors for predicting task events (see table 1, **Task** row, **MMI** columns), we extracted 3 second crops as in prior work [36] starting from the event offset of sessions 4, 8 and 12 where the subjects imagined opening and closing their left or right fists (left or right being the target class). Unless otherwise stated, the EEG channels were limited to those in common with the BCIC dataset.

**Table 1.** Accuracy of predicting targets over five-fold cross-validation using majority label of five nearest-neighbours. Only the prediction of subject and dataset scored appreciably over chance (all but the dataset target were balanced; chance level prediction was 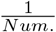). Sensitivity to variation in T-Vectors was accounted for by comparing single T-Vectors through an average of four sequential vectors. This averaging showed a consistent, though mild trend in subject prediction. The dashed MMI/Dataset row is because this is the same experiment as BCIC/Dataset (and has the same uniform 100% prediction).

|  |  | BCIC |  |  |  |  | MMI |  |  |  |
| --- | --- | --- | --- | --- | --- | --- | --- | --- | --- | --- |
| Target | Num. | 1 | 2 | 3 | 4 | Num. | 1 | 2 | 3 | 4 |
| Subject | 9 | 89.2 | 93.5 | 95.5 | 96.1 | 105 | 92.2 | 96.7 | 97.6 | 97.7 |
| Session | 2 | 53.2 | 52.5 | 52.6 | 50.8 | 14 | 6.7 | 6.5 | 6.6 | 6 |
| Task | 4 | 27.1 | 28.0 | 26.7 | 26.3 | 2(L/R) | 53.2 | 54.5 | 55.3 | 53.5 |
| Dataset | 2 | 100 | 100 | 100 | 100 | — | — | — | — | — |

## Methods

### Training Procedure

Datasets in DN3 provide a mechanism for accessing subject IDs paired with each fetched point. These are used as the labels and inputs respectively for pretraining the T-Vector network with the TUEG data. Each point was linearly scaled and shifted so that its largest value was 1 and smallest value was −1 (note this was *not per channel*, and the entire point/trial was scaled and shifted by the same factors), which was also done when extracting T-Vectors of downstream data.

Here, we simply employed the network used to create X-vectors [37], but reduced the hidden size of the network to 384 rather than 512. Each T-Vector is then the 384-element-long hidden representation from layer `segment6` (as labelled by the original authors [37]). These are the activations of the second to last layer (third to last if including the final `softmax`) before the non-linearity. The entire network was of course subject to training during the pre-training stage, with the final `softmax` creating a distribution over the 1364 targets (people).

Optimization was performed in batches of 128 using the ever-popular Adam optimizer [38] with a default learning rate of 0.001. This rate was divided by 10 at epochs 50 and 75 through the 100 epochs of training. A minimal L2 weight-decay was added to the loss at a factor of 0.00001 (larger values appeared to not separate clusters as well during tuning with a small subset of 100 people).

To minimize sensitivity to any *expected* length, we further cropped each loaded *batch* uniformly to 20%-100% of its original length. Thus the network is trained to identify users using as little as a single second’s worth of data, with no particular consistency in task. This strategy is implemented as a DN3 batch transform, and is part of the available transforms in the library.

At the end of pre-training, the model weights were frozen and no longer updated. The final weights used can be downloaded here and DN3 provides tools to easily recreate the network with these weights. Finally, the T-Vector representations of each point of each downstream dataset was collected and saved for analysis.

### Analysis of vectors

We analyze T-Vectors with two complementary approaches. First, we consider a simple supervised prediction of notable variables using *k*-nearest-neighbours (*k* = 5 is used throughout). Naturally, subject identity was the most critical variable considered, but we also included: session identity, which task was being performed (e.g., canonical trial labels such as left versus right hand motor imagery task), and dataset prediction (mixing T-Vectors from both downstream SMR datasets). The additional variables were to highlight if the spatial distribution of the points was informative of any other known quantity besides subject identity. These were all compared using 5-fold cross-validation, stratified by prediction target. That is, each fold had an equal (as possible) percentage of each target variable. Our second analysis visualizes the manifold of T-Vectors using t-distributed stochastic neighbour embeddings (t-SNE), noting how readily separable the T-Vector space appeared, and if there were any ready interpretations of behaviour outside of this. Throughout, we set the perplexity of the t-SNE operations to 30.

Throughout, we additionally consider the effect of smoothing the single trial T-Vectors by averaging up to 4 sequential vectors to minimize T-Vector variance. This averaging never crossed the boundaries of covariates considered (i.e., vectors from one session and a second session were not averaged together when predicting session); instead, the final point would simply be an average of the remaining points.

## Results

### Variable prediction

Table 1 clearly demonstrates that the local (Euclidean distance) space around each T-Vector is highly informative of subject identity. Predicting vectors from the held-out fold using the closest vectors of the remaining folds yielded results that were, in all but one case, well over 90% accurate at identifying subject (and the remaining case was very near this point). Conversely, the local space of the T-Vector representation was markedly less informative of which session or task was being performed, irrespective of dataset. Interestingly, identifying which dataset T-Vectors belonged to was profoundly accurate, making no mistakes. Noting that this variable can be seen as a mixture of subject variation and some other dataset-specific variation, it is clear that some information besides subject identity is encoded by the T-Vector representation. At the very least, whatever confused the subject identity prediction *within* a dataset did not extend *between* the datasets considered.

### Manifold visualizations

We observe a general tendency for points from the same subject to have a common localization in Figure 3, becoming stronger after pooling. This suggests that the vector representation is robust under different conditions. The BCIC dataset allowed for observing any variation across sessions separated by days (in figure 3, star and cross points represent the different sessions). It remained clear that, for the most part, different sessions did *not* separate into different clusters. Showing some stability over time. Subject A05 was a notable exception to this (and is most obvious after pooling; purple points in Figure 3b). Furthermore, there was a notable region of ambiguity in Figure 3a made up of points from subjects A01, A02, A03, and A05 (cluster made predominantly of blue, orange, green and purple respectfully, although other colours also added to the mix). While patterns like this are hard to interpret using a single t-SNE plot (or even several for that matter), a reasonable correlation between this zone and subject-specific performance was also observed. The subjects where we observed the lowest single-vector (subject) classification were also A01, A02, A03, and A05 – all of whom scored below 90% (with the remaining subjects scoring above this mark). The supplementary table in Appendix B provides more subject-specific details. After considering the subject-specific performance in conjunction with 3, we concluded that the visualization is representative of how T-Vectors separate data from unseen subjects performing unseen tasks, with novel hardware. In other words, T-Vectors do seem to generalize to new datasets and subjects without any further adaptation.

**Fig 3.**
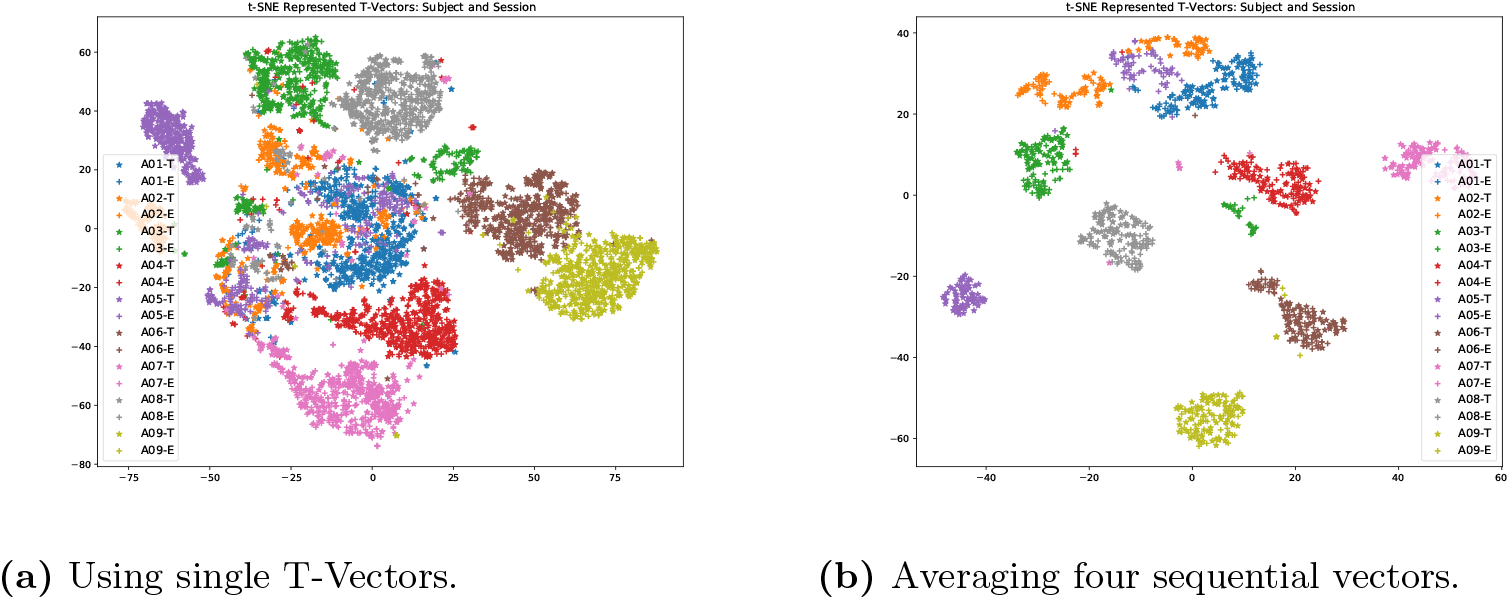
t-SNE visualizations of the high-dimensional manifold of the learned T-Vector space. The nine subjects of the BCIC dataset are plotted in different colours, while their separate training and evaluation recording sessions are marked with stars and crosses respectively. For the most part each subject’s vectors were separately clustered, though subjects A01 and A02 were notably less clustered. Averaging across four vectors in (b) dramatically reduced cluster overlap, but not entirely.

This pattern of subject separability is all the more clear in Figure 4, in which the two colours represent the two downstream datasets. Rather than isolating two major groupings (one for each dataset), the pattern indicates sets of localized structures correlated to subject identity. Many small pockets of data abound and, after counting all groupings that did not consist of individual points, the number of MMI clusters is either 104 or 105 (the center region has some ambiguity), which corresponds to the 105 subjects used from this dataset (recalling the high accuracy from Table 1, these groupings were largely homogenous). The BCIC groupings appear to provide approximately 8 groupings, one fewer than the total subjects, but with a distinct cluster reminiscent of the most ambiguous region of Figure 3b, although this is not conclusive. Very little changed when adjusting for the differences in recorded channels, focusing only on the 22 EEG channels common to both datasets, Figure 4a looks largely identical to Figure 4b albeit, in the former, the MMI dataset appeared to encircle the BCIC dataset. We therefore conclude that, while determining the source dataset of a particular T-Vector is readily apparent from its neighbours (see Table 1), this was (for the most part) a consequence of separating the independent groups of subjects. In other words, a stability of subject-wise representation is shown across datasets which were recorded very differently.

**Fig 4.**
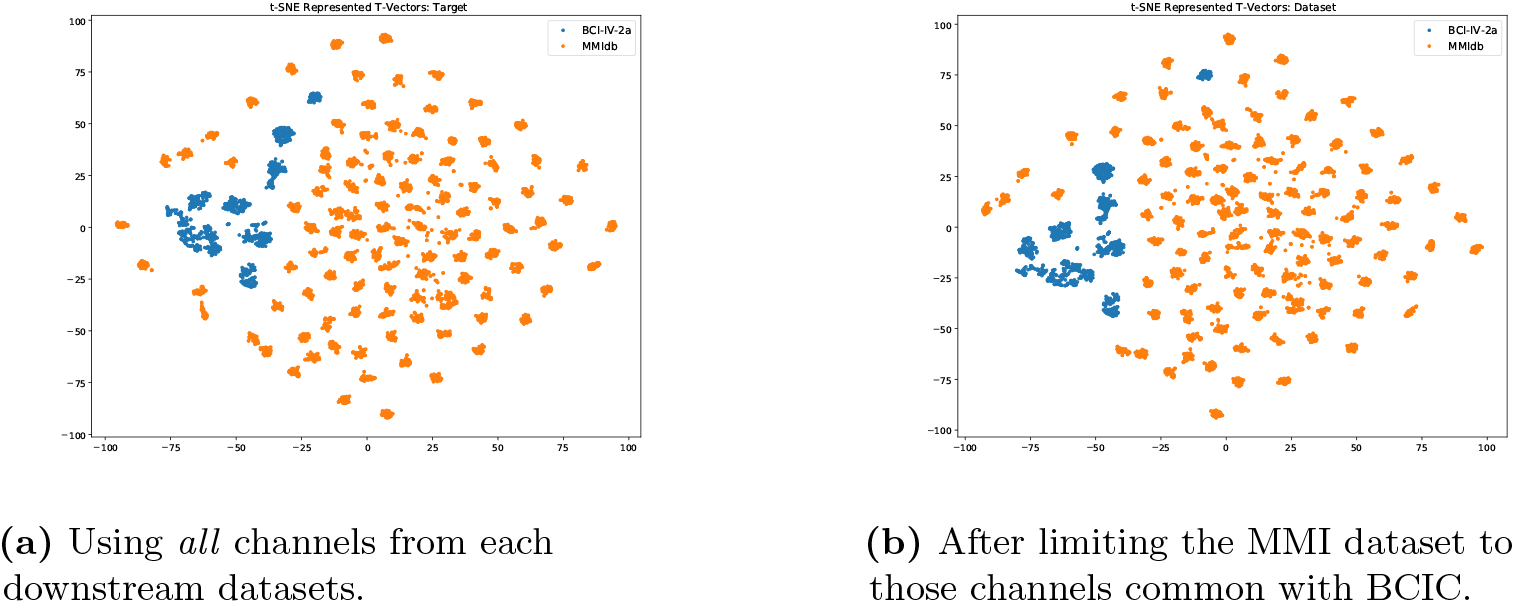
t-SNE visualizations for T-Vectors across both BCIC and MMI datasets (blue and orange respectively). The two figures show the difference in distribution when in (a) excessive channels were used with the MMI dataset: including additional channels not present in the pretraining set, and (b) limiting to the mostly 10/20 channel set common to both. The figures are largely identical, indicating that the extra channels were ignored. The major commonality between (a) and (b) is that each subject has roughly one cluster to account for them, with minor ambiguity. By visual inspection, we noted 8 representative clusters for 9 subjects for BCIC and 104, perhaps 105 clusters out of 105 subjects for MMI.

### Discussion of T-Vectors

T-Vectors are promising for identifying individuals using minimal EEG recordings. While these vectors may be effective as presented, we expect that without introducing some notions of fairness [39], biases are likely to occur. For example, we might observe a greater sensitivity for well-represented demographic intersections in the data for better represented groups, than for other demographics.

We also warn against quickly interpreting T-Vectors (or features therein) as strongly correlated to intersections of demographics or notions of personal characteristics (e.g., intelligence). It would be an error to describe T-Vectors as an “objective” representation of variability – instead, they simply capture some latent features that seem to disambiguate individuals well. Specific investigation into any correlations needs to be done subsequently, with consideration of the likely data biases mentioned above.

With these considerations in mind, we intend to explore possible correlations between T-Vector components and markers of mental health, in addition to better qualifying how sensitive T-Vectors are to sessions separated by longer time scales, more different hardware, recording modalities (e.g. transfer to magnetoencephalography) and performed tasks. While capturing correlations like this is certainly a potentially interesting avenue, T-Vectors may also be informative of the scale of these differences, e.g. by distance along a direction of variation, a somewhat unique aspect of this continuous latent space approach.

As mentioned in our motivation, prior work in transfer learning has considered the use of **adaptive classifiers**, whose major mechanism of performance transfer comes from identifying which users are most *similar* ^13^. Future work should consider if T-Vector distance can be used towards similarly identifying like-users. Again as mentioned above, previous work has found *alignment* of subject’s data along the SPD manifold to be an effective method for creating general classifiers. Future work will consider if a data transformation scheme can do the same in T-Vector space, with the advantage of the T-Vector space being more ready use of Euclidean notions of distance rather than Riemannian.

Additionally, throughout all t-SNE visualizations there are a variety of singular points scattered throughout. While these were minimized after averaging sequential vectors (see figure 3b), they are never removed entirely. It would be prudent to consider what these outliers are if they remain after further development. Could they for instance be overly contaminated with muscular artifacts, or other notable characteristics? In this way, it is worth considering if T-Vectors may also prove to be a quick form of data triage, detecting usable versus non-usable data given an expected template T-Vector.

In terms of biometric applications, the performance levels presented above are similar to previous work. For example, recent work with the MMI dataset was over 99% accurate, with a reduced channel set and shorter time window [40]. However, this result is achieved by training a DNN with the first 90% of each session’s data, and predicting the identity of the user with the remaining data. While this is not uncommon in the literature [30], this paradigm does not necessarily generalize across datasets or hardware and may introduce channel effects and even information leakage which artificially boosts performance. We are unaware of any prior work that does in fact generalize in this fashion, but it is clear that this is a desired property for the application [30]. We therefore suggest that T-Vectors represent the state-of-the-art despite not reporting the greatest performance. Towards resolving a claim like this, previous work has considered the development of a score for EEG biometrics [31] called a *U* score, but we find some difficulty in fairly calculating *U* for our own work, as no other prior work performed such a great degree of pre-training to develop their features in the first place. Specifically, *U* rightfully aims to minimize the amount of time needed to identify a subject, which is determined in terms of total time of recordings for training. It is ambiguous how *pre-training* would be factored in. If it is, by virtue of this large number our method has an inconsequential score. If we focus only on time to develop a single prediction, i.e. the time embodied by the nearest-neighbours used for prediction, our own method outperforms the previous best score by 3.5%. However, in the interest of not excluding pre-training approaches, and further extending the score to evaluate generalization across *multiple* datasets, we propose some revisions in appendix C.

The source-code for this project can be found at https://github.com/SPOClab-ca/T-Vectors and can be seen as a template for other DN3-based projects. The particular T-Vector weights used in these analyses can be downloaded here.

## Conclusion

We have presented DN3, a new Python-based deep learning library and set of APIs for BCI and more general neuroscience applications. DN3 aims to increase reproducibility while minimizing redundancy at little to no expense in flexibility. Furthermore, we aim it to be a community-driven solution to engage with DL techniques that might otherwise be inaccessible. Intended additions to DN3 include more applications of large datasets such as semi- and self-supervision tasks, meta-learning optimization and adversarial learning processes. All the while, we hope to continue to integrate more multi-purpose preprocessing steps, transforms, and trainable modules as they develop. These goals ultimately can only be confirmed by the community at large, but we have presented a unique application of DL with EEG data that was considerably streamlined through the use of DN3.

## Acknowledgments

We would like to emphasize our thanks to those who make data they have recorded publicly accessible to all. Rudzicz is supported by a CIFAR Chair in Artificial Intelligence. Thanks to Sicong Huang for revitalising an interest in the T-Vectors idea.

## A TUEG dataset triage

Unlike the MMI and BCIC datasets, the TUEG dataset had little if any inspection for data quality. Furthermore, the number of channels can vary, in some instances files indicate that a channel was recorded, but it ends up blank. As this scale of data is profoundly useful, but difficult to visually inspect for artifacts, errors and the like, we considered the difference in distribution of an easily acquired statistic: the standard deviation of channel values for each trial, to narrow the focus on which trials were likely best representing *good* data.

Figure 5 shows that the TUEG and MMI distributions are quite distinct. While it is hard to conclude what *good* data should look like, this plot clearly shows where TUEG exhibits some *bad* features. The largest peak of the histogram is a relatively narrow section localized at (and very near to) a standard deviation of 0. Such points are not representative of any real EEG data and needed to be excluded. Additionally, given that some of the TUEG dataset comes from clinical evaluation of epilepsy patients, and clinical settings in general that may have floating or unconnected channels, the large representation of points with standard deviations greater than 0.6 were suspect to us when compared to the MMI distribution. To focus on data that seemed somewhat representative of our ultimate application, we specified upper and lower cutoffs in the range that seemed to (by inspection) match the MMI distribution. Thus, points with channel standard deviations *σ*_*trial*_ were only used in training if 0.04 ≤ *σ*_*trial*_ ≤ 0.45.

**Fig 5.**
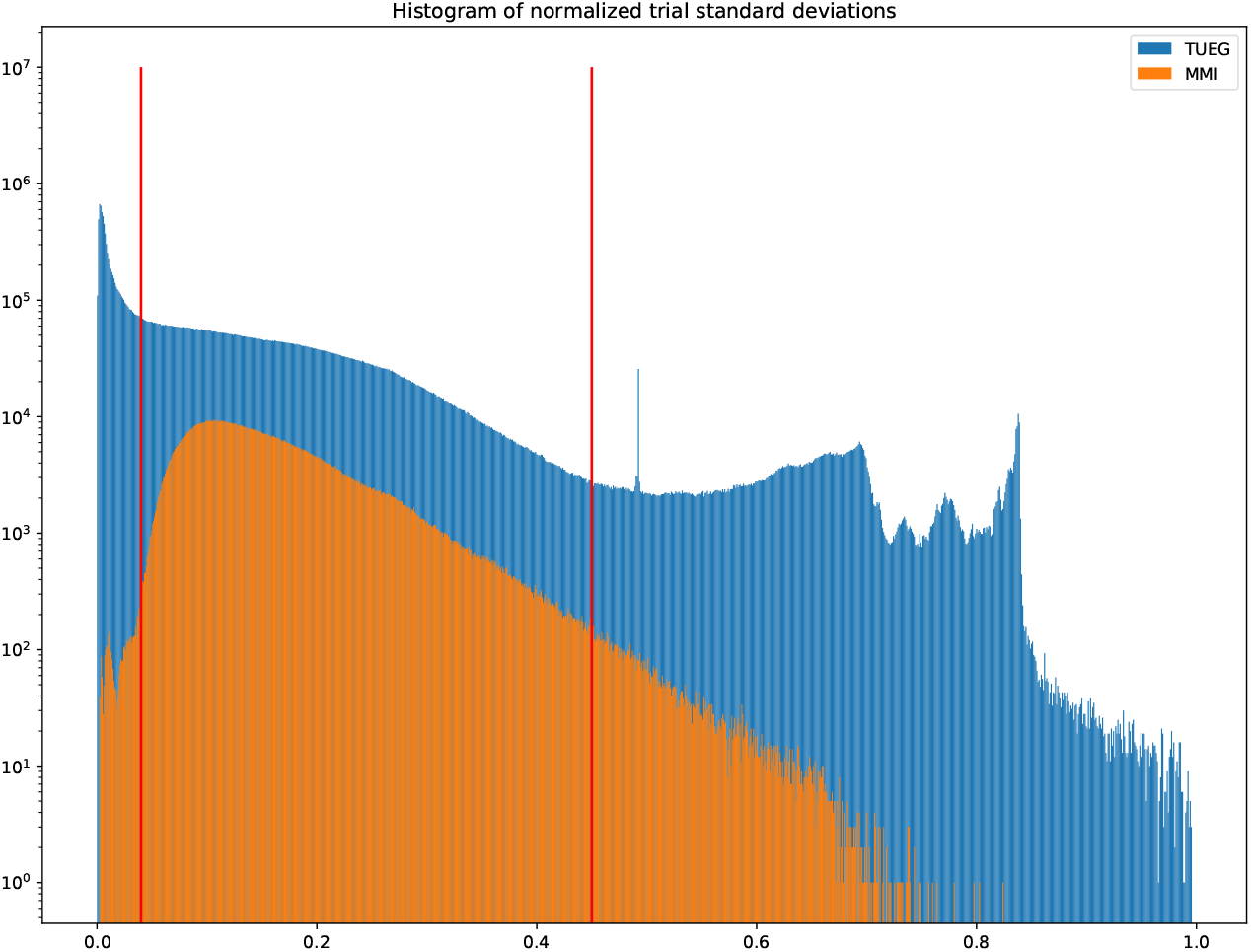
Log-histogram comparing the distribution of channel-wise standard deviations for all normalized points (cropped training sequences) in the MMI and TUEG datasets. Notice the large peak for TUEG at a standard deviation of 0, and the surprising (in relation to the MMI dataset) increase in deviation ≥ 0.8. We chose to narrow the selection of usable TUEG trials to those that were between the red lines (0.04 ≤ σ_*trial*_ ≤ 0.45), simply so that the noted extremes were rejected.

## B BCIC subject-specific performances

**Table 2.** Accuracy according to each subject across all five folds of data for the BCIC dataset. Each row considers how many sequential vectors were averaged to develop points. Notably, A01 underperforms other subjects, but averaging sequential vectors compensates for this problem. Almost uniformly, this averaging improves performance for all subjects.

| Subject | A01 | A02 | A03 | A04 | A05 | A06 | A07 | A08 | A09 |
| --- | --- | --- | --- | --- | --- | --- | --- | --- | --- |
| 1 | 73.0 | 84.9 | 89.0 | 92.4 | 87.8 | 92.2 | 98.5 | 92.0 | 97.3 |
| 2 | 83.1 | 91.9 | 94.1 | 95.5 | 87.5 | 94.6 | 99.6 | 97.2 | 100.0 |
| 3 | 90.3 | 95.2 | 94.8 | 95.9 | 91.0 | 95.9 | 100.0 | 97.5 | 100.0 |
| 4 | 93.3 | 95.2 | 96.3 | 95.2 | 93.2 | 95.3 | 100.0 | 96.6 | 100.0 |

## C Subject identification score

In the survey performed by Yang & Deravi on using EEG for biometrics [31], the authors proposed a formulation of a general score to compare subject identification across a range of contexts and datasets. This score was designed to increase in proportion to the number of subjects *N* (train and test since prior work developed capacity to identify a fixed set) and the overall subject identification accuracy *Acc*. This was balanced against minimizing the length of time needed to identify a subject *T*_*e*_ and the number of channels used *C* to do so. All while also minimizing the amount of data needed to train such a system *T*_*r*_.

Their overall score *U* was then:

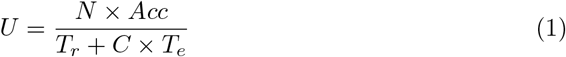

The highest overall score they ultimately determined was 86.63 scored while similarly using the MMI dataset [31].

Considering whether our own work could be scored in this fashion, it is immediately clear that the *T_r_* term (specified in seconds) would be exceedingly large (1.6 million seconds) and the score would be near zero. If however, we were to take some liberty with the interpretation of this quantity, and identify it as the amount of time needed to identify a *test point*, one could plausibly see this term as the time embodied by the five neighbours employed in our nearest-neighbours classifier above. Thus, we might say *T_r_* = 5 4*s* = 20*s*. The upper bound on the score for MMI identification with T-Vectors as presented in the main body of this article would then be:

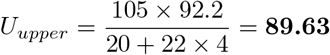

While this is higher than the previous best, this is to be sure, an *upper bound*. That being said, we would like to propose an alternative score that is more amenable to methods that:

1. Can be uniformly leveraged over multiple datasets/hardware
2. Can employ whatever means effective to *develop* features
3. Can support different channel, reference and evaluation quantities with respect to each person and trial

To this end, we propose to re-consider the *T*_*r*_ term so that it is more clearly defined as: *the amount of time (in seconds) needed to develop a prediction with accuracy* 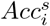 *of a particular point (sequence)* 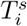. Where both terms are with respect to a point *i* ∈ *I*_*s*_, where *I*_*s*_ are all the available *testing* points for subject *s* ∈ *N*_*subjects*_. In other words, *T*_*r*_ for the T-Vectors methodology is the sum of time employed for the k (in our case 5) neighbours needed to develop a prediction. Considering a DNN trained to simply identify *N*_*subjects*_, this 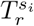 would be the total time points used in the training data for that particular subject. Furthermore, 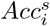 then is a binary variable, where the value is either 0 or 100, indicating whether each point is predicted. The choice of 100 rather than 1 is simply so that the resulting score will tend to be greater than 1, and is mostly compatible with the prior work. Additionally, we suggest that the score should be *increased* in a reasonable manor with respect to the number of different datasets *N*_*ds*_ (ideally hardware) that performance carries over for. The full expression of our revised score is then:

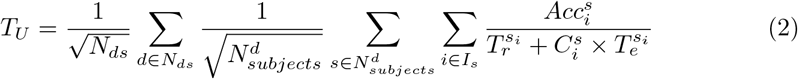

Fundamentally, this is a revision of the *U* score, as at its core it remains mostly the same. In fact, *U*_*upper*_ is nearly recoverable from this general formulation, since we calculate it using a single dataset, and *T*_*s*_, *T*_*c*_ and *C* parameters that do not vary in subject or reference trial. The only difference in score here would be the division by the square root of the number of subjects (and datasets), which we added to minimize simply surpassing the state-of-the-art by adding inconsequential additional subjects or datasets. This rather should reflect larger demonstrations of generalization across people and datasets.

While we expect that adding any other dataset using the pre-trained T-Vectors would quickly overtake this score, the *T*_*U*_ score of the work in the main body of this article would be:

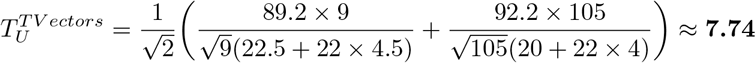

## Footnotes

1 https://en.wikipedia.org/wiki/MNIST_database

2 https://pytorch.org/

3 https://dn3.readthedocs.io/en/latest/

4 https://www.mathworks.com/products/matlab.html

5 The most common being a variation used by datasets found at physionet.org, which holds a number of commonly used public datasets. Additionally, recordings and subjects can be identified using a simple pattern-matching scheme.

6 Note that as described

7 Currently using the whole project, which includes other versions of the data results in over ten thousand targets and has proven more challenging and will be considered in future work.

8 See the uploaded configuration file for more details.

9 https://www.isip.piconepress.com/projects/tuh_eeg/html/downloads.shtml access requires a simple sign-up process.

10 https://physionet.org/content/eegmmidb/1.0.0/

11 Originally here http://www.bbci.de/competition/iv/, which requires request for access, but we found some difficulty with these files and instead converted the unrestricted copy found at http://bnci-horizon-2020.eu/database/data-sets to the native MNE (raw.fif) format.

12 This is also a default aspect of the Deep1010 mapping. Additional default behaviour includes mapping one of the auxiliary channels to a global scale parameter, which represents the factor by which a particular point’s maximum absolute value (the 1 or −1 value in the point) relates to the absolute maximum value in the dataset, as specified in the configuration file. This allows for consistently scaled trials, while still informing models as to the scale of the point’s context. This was not used for T-Vectors to minimize any average amplitude being suggestive of identity.

13 Note that t-SNE plots do not represent distance well beyond local structure and in fact can be misleading when referring to the “closeness” of different locally coherent areas, we caution against using the plots above for this purpose.

